# Measures of Implicit and Explicit Adaptation Do Not Linearly Add

**DOI:** 10.1101/2022.06.07.495044

**Authors:** Bernard Marius ’t Hart, Urooj Taqvi, Raphael Q. Gastrock, Jennifer E. Ruttle, Shanaathanan Modchalingam, Denise Y.P. Henriques

## Abstract

Moving effectively is essential for any animal. Thus, many different kinds of brain processes likely contribute to learning and adapting movement. How these contributions are combined is unknown. Nevertheless, the field of motor adaptation has been working under the assumption that measures of explicit and implicit motor adaptation can simply be added in total adaptation. While this has been tested, we show that these tests were insufficient. We put this additivity assumption to the test in various ways, and find that measures of implicit and explicit adaptation are not additive. This means that future studies should measure both implicit and explicit adaptation directly. It also challenges us to disentangle how various motor adaptation processes do combine when producing movements, and may have implications for our understanding of other kinds of learning as well. (data and code: https://osf.io/dh86e)

## Introduction

Both implicit (unconscious, automatic) and explicit (conscious, intentional) processes contribute to various kinds of learning (Jacoby, 1991). Research exploring the contribution of implicit and explicit processes to human motor adaptation relies on the notion that these processes are related (Benson et al., 2011; Taylor and Ivry, 2011; Werner et al., 2015). Many recent studies assume they linearly add to total adaptation, with some support (Bond and Taylor, 2015; Redding and Wallace, 1993; Sülzenbrück and Heuer, 2009). Here, we test whether this additivity assumption holds.

The main idea is that there are only two kinds of processes contributing to adaptation: those we are aware of, and those we are not aware of. The processes we are aware of, which are usually intentional, involve a strategy, can be verbalized, and are often effortful, are referred to as “explicit” adaptation. The processes we are not aware of are then not intentional but automatic, they can not be verbalized and since they are automatic they usually require less effort, and these kinds of processes are often called “implicit” adaptation. A further distinction is that explicit processes can be voluntarily disengaged, as opposed to implicit processes. Implicit and explicit adaptation could each be split up further, or overlap with other kinds of processes (e.g. reward-based learning could be both implicit and explicit). However, if we can split motor adaptation into implicit and explicit processes, it seems intuitive that we can also add them for an estimate of total adaptation, the “additivity assumption” (Maresch et al., 2021a):

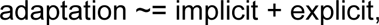

Often illustrated as in Figure 1A. This relationship, if true, could be rewritten as:

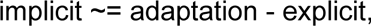

**Fig 1:**
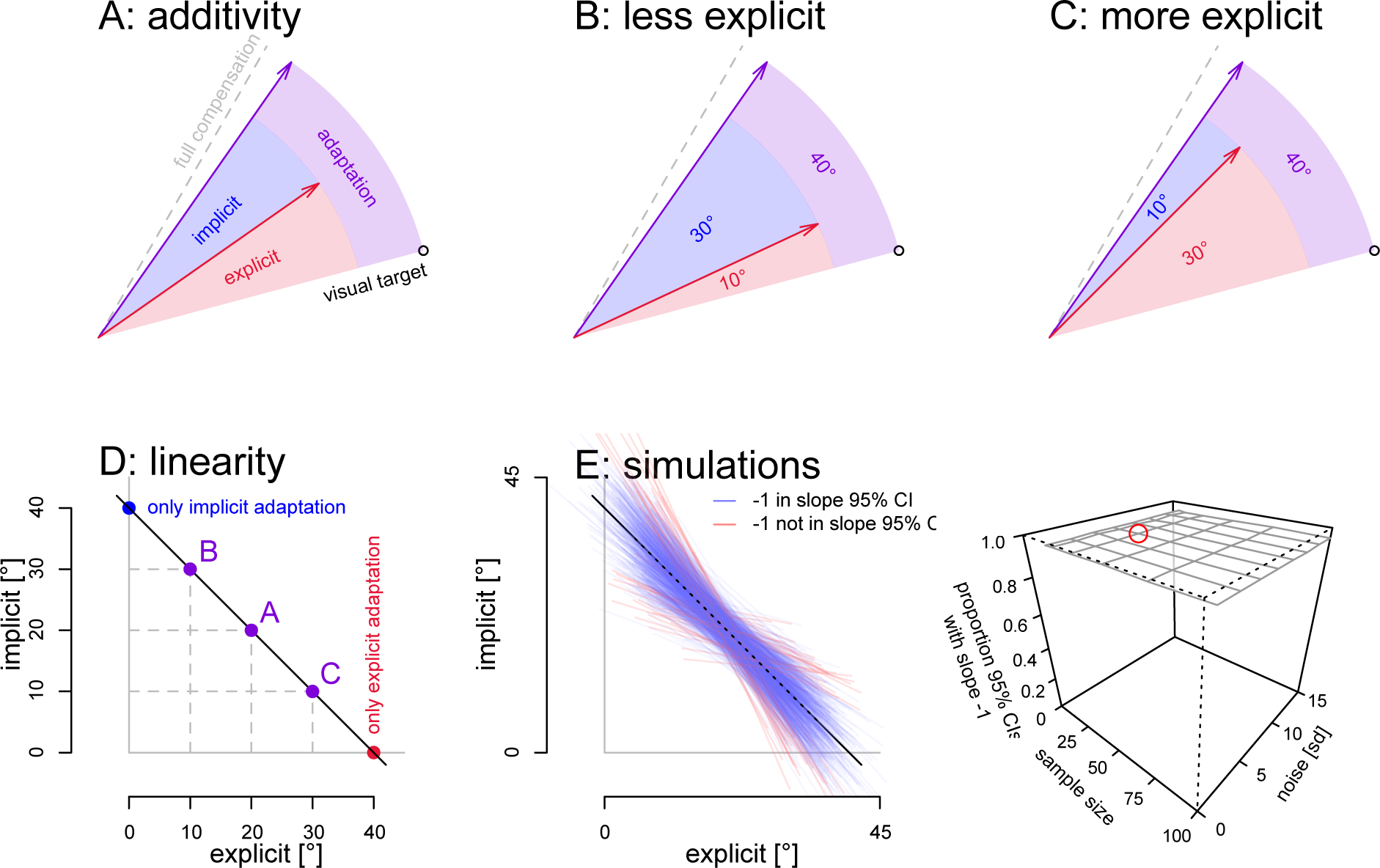
The additivity assumption. **A:** Often, implicit adaptation (blue) is considered what is left after explicit adaptation (red) is subtracted from total adaptation (purple). **B,C:** Given less explicit adaptation, this would mean there is more implicit adaptation (B), and vice versa (C). **D:** This means that if explicit and implicit adaptation are additive, they should have a negative, linear relationship. The purple points depict the situations in panels A,B,C and also shown are situations where adaptation is fully implicit (blue) and fully explicit (red). **E:** We generated 1k data sets (sample size: 24) with the mechanism prescribed by the additivity assumption adding normally distributed noise (sd: 7), and test if we can recover the predicted negative slope from a linear model fitted to each data set (left). In ∼95% of simulated data sets, the 95% confidence interval of the slope includes the predicted value of -1 (blue lines) and in ∼5% it doesn’t (red lines). Finally, we test different sample sizes and levels of noise, but that does not affect the results of our test: in all cases ∼95% of the simulated data sets the 95% CI for the slope includes -1 (right). Red circle: the combination of sample size and noise used on the left.

In Figure 1A, implicit adaptation (blue wedge) is depicted as what is left when explicit adaptation (red arrow) is subtracted from total adaptation (purple arrow). In theory, the additivity assumption allows researchers to determine the implicit contributions to motor adaptation without the need to directly measure implicit learning. In practice however, the underlying assumption is not always replicated (Modchalingam et al., 2019) and has been called the “least robust” assumption in motor adaptation (Maresch et al., 2021a).

Some of the papers our lab published in the last five years (Gastrock et al., 2020; Modchalingam et al., 2019; Vachon et al., 2020) used the Process Dissociation Procedure (Jacoby, 1991; Werner et al., 2015) to assess implicit as well as explicit adaptation. In the first paper we already noticed that reach deviations in include strategy no-cursor trials did not equal the reach deviations during training trials. That is, the difference between include and exclude no-cursor reach deviations would not necessarily equal the magnitude of explicit strategies. Hence, in the two later papers we just took the existence of any difference as definite evidence for an explicit strategy. This also lead us to question the additivity of explicit and implicit adaptation. In particular, it seemed to us that there are two very separate neural processes that likely manifest in different brain areas or even networks. It would be very unlikely for such different processes to linearly add - that would simply not be a biologically plausible mechanism. It may also have to do with the nature of the measurements used in the Process Dissociation Procedure, but these do have one distinct advantage: there are 2 no-cursor measures as well as adaptation. Using these 3 different measures it is possible to notice any kind of discrepancy resulting from non-additivity. Nevertheless, we should not have relied on additivity and our PDP style approach to assess explicit adaptation.

In recent motor learning work, explicit adaptation is instead measured as re-aiming reports. While this provides a better measure of explicit adaptation, it is often combined with the additivity assumption to forego measuring implicit adaptation. Instead, implicit adaptation is taken as the difference between total adaptation and explicit adaptation. A substantial portion of findings rely on such “indirect” means to determine implicit adaptation (McDougle et al., 2015; Miyamoto et al., 2020; Wilterson and Taylor, 2021).

Both the PDP-style approach as well as re-aiming based approaches on their own rely heavily on the additivity assumption as they either have no direct measure of explicit adaptation or no direct measure of implicit adaptation. In a set of experiments we attempt to validate if the assumption of additivity would allow for the “indirect” method of determining implicit or explicit learning to be equivalent to direct measures. In particular, the field seems to focus on using reaiming measures and subtracting them from total adaptation as a measure of implicit learning. So this is where we will focus as well, although for completeness we also test additivity in data collected with a PDP approach.

The statistical approach we follow relies on the negative correlation between implicit and explicit adaptation that would be dictated by additivity. That is, when total adaptation is constant, and explicit adaptation decreases, implicit adaptation has to increase by the same amount (Fig 1B) and vice versa, when explicit adaptation increases, implicit adaptation decreases (Fig 1C). That relationship between implicit and explicit adaptation under additivity can be plotted as a linear function with an intercept equal to total adaptation and a slope of -1 (Fig 1D). To see if this would be reproducible in principle, we ran a simulation where we added noise to data generated with an additive model. All noise (ε) was drawn separately from a normal distribution with μ=0° and σ=7° (results are independent of the level of noise). For each participant *p*, total adaptation and explicit adaptation are given by:

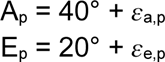

And implicit adaptation is taken as the difference between total adaptation and explicit adaptation (plus noise):

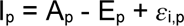

We can repeat this for a number of participants and then fit a linear model predicting the generated implicit adaptation from the corresponding explicit adaptation. We will see how well a linear model fitted to the generated data recovers the slope of -1. (The intercept should be close to total adaptation, but since most of the data should be somewhat removed from zero explicit adaptation, the intercepts will vary much more and this might not be as good a test.) In particular we can test if the 95% confidence interval of the mean for the slope parameter includes -1. Using samples of 24 participants (equal to our experiment, see below) we’ve repeated this 1000 times (linear fits shown in Fig 1E, left), and in ∼95% of simulations the 95% confidence interval of the slope includes -1 (blue lines in Fig 1E; red lines: 95% CI for the slope excludes -1). Finally, with 10k simulations each for various combinations of sample size and noise, we show that this statistic is independent of the level of noise or chosen sample size (Fig 1E, right) as would be expected with normally distributed noise. That is, in data that is generated according to additivity (plus noise), the linear relationship between implicit and explicit adaptation with a slope of around -1, can be recovered from noisy data. This means that the 95% confidence interval of the slope can be used as a statistical test, with alpha=0.05, of additivity of implicit and explicit adaptation.

The simulations with a generative model shown above, rely on the same linear relationship between implicit and explicit adaptation as previous tests of additivity (Bond and Taylor, 2015; Redding and Wallace, 1993; Sülzenbrück and Heuer, 2009). However, previous tests only looked for significant correlations. That is; any non-flat linear relationship was taken as evidence for additivity. Here we test an additional property of data generated under additivity: a particular slope. There could be additional relevant properties (e.g. based on the intercept or residual errors) but the confidence interval for the slope of the linear relationship already provides a stricter test. We will use this approach and variations on it in several tests of the additivity assumption, both in data from our own experiment and in an aggregate data set with data from several previous papers representing various labs, setups and paradigms.

### Varying explicit adaptation in strict and loose additivity

If total adaptation is the sum of implicit and explicit adaptation, then, given (near) constant total adaptation, implicit and explicit adaptation should perfectly complement each other. That is: under the additivity assumption, a given change in explicit adaptation should result in a change in implicit adaptation that is of equal size, but in the opposite direction. And this should also be true across participants: a participant with higher explicit adaptation should have lower implicit adaptation and vice versa (see Fig 1). Thus, we did an experiment with three conditions that we expected to result in different levels of explicit adaptation, but highly similar total adaptation. The main condition has independent measures of implicit and explicit adaptation, the other two set the context.

## Methods

### Participants

For this study 72 right-handed participants (55 female; mean age 19.8, sd 3.7; demographics not stored for 1 participant) were recruited from an undergraduate participant pool, 24 for each of 3 groups (aiming, instructed, control). All participants reported having normal, or corrected-to-normal vision and gave prior, written, informed consent. All procedures were approved by York University’s Human Participant Review Committee.

### Setup

The participants made reaching movements with a stylus on a digitizing tablet, controlling a cursor that was displayed on a downward facing monitor (60 Hz, 20”, 1680×1050, Dell E2009Wt) seen through an upward facing mirror in between the tablet and monitor (see Fig 2A). This puts the perceived stimuli in the same plane of depth as the hand, and the displacement of the cursor was scaled to match the displacement of the stylus.

**Fig 2:**
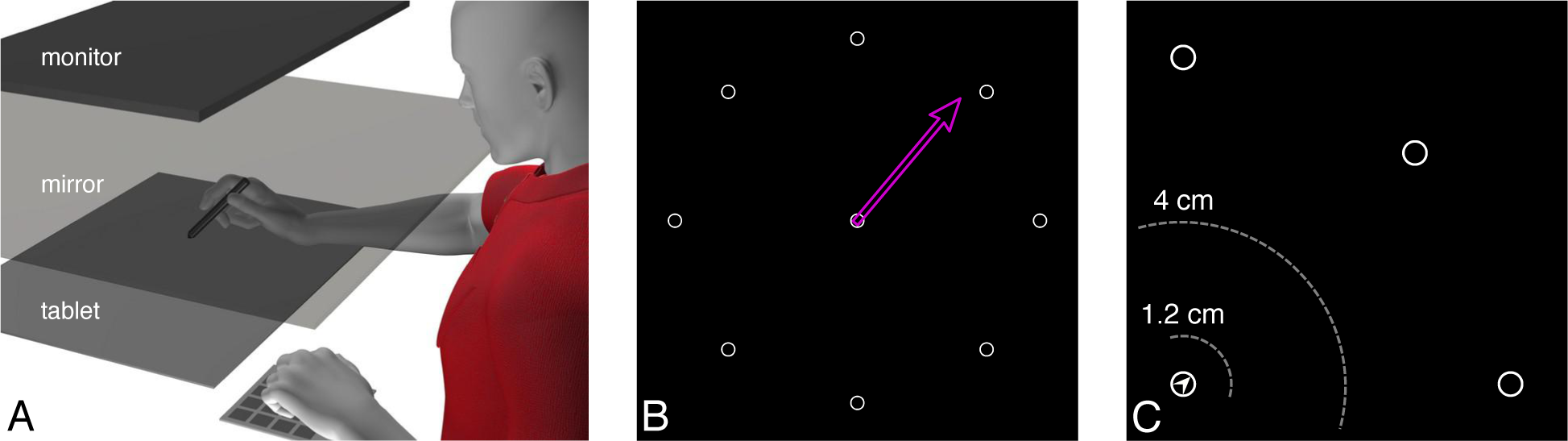
Setup, aiming and no-cursor trials. **A:** Tablet setup with mirror. The stimuli appear in the same plane of depth as the tablet. **B:** Aiming: the upcoming target (1 of 8) is shown with an arrow used to indicate the intended reach direction, or “aim”. **C:** The outward reach on no-cursor trials ends when the stylus stops moving for 250 ms and is at least 4 cm away from the home position (outer gray ring). For the return home, a small arrow-head in the home position indicates at 45° intervals where the cursor is, and participants move in the opposite direction. When the stylus is then back within 1.2 cm distance of the home position, a cursor appears that needs to be moved to the home position to start the next trial.

### Stimuli

The background of the screen was kept black. An open gray circle (radius: 0.25 cm) served as a home position and was located at the center of the screen (see Fig 2B). Eight reach targets were located at 8 cm from the start position at 0°, 45°, 90°, 135°, 180°, 225°, 270° and 305° and were also displayed as open gray circles (radius: 0.25 cm). A filled circle was used as a cursor (white disc, radius: 0.25 cm) on training trials. In the aiming group only (see below), an open pink arrow originating at the home position and extending 7 cm was used on aiming trials (it initially deviated ±10° or ±5° from the target, randomly chosen on each trial). On no-cursor trials a small gray arrow head was presented within the home position, pointing to where the unseen cursor would be, in intervals of 45°, to guide participants back to the home position. More details below.

### Tasks

In each trial, participants used the stylus to move from a central home position to one of the 8 targets. Before each reach, participants have to keep the cursor at the home position for 2.0 s (trials without aiming) or 0.5 s (trials with aiming). After the hold period the target would appear and the home position would disappear, signaling to participants they should move to the target

### Training trials

In training trials, a cursor had to be moved from the home position to the target and back. When the cursor’s center was within the radius of the target, the reach was considered ended. At that point the target disappeared and the home position reappeared, signaling that the participant should move back to the home position. Once the cursor’s center was within the radius of the home position, the trial ended. During the aligned phase, the cursor was aligned with the tip of the stylus. In the rotated phase, it was rotated 30° around the home position.

### No-cursor trials

In no-cursor trials, the cursor disappears as soon as it’s center no longer overlaps with the home position. The reach is considered ended if the stylus has moved beyond 50% of the home-target distance (4 cm, Fig2C) and there is no movement for 250 ms (or the total movement during the last 250 ms is less than 1% of the home-target distance). At this point, the target disappears, the home position reappears, and an arrow at the home position indicates where the (rotated) cursor would be relative to the home position (in increments of 45 degrees). Participants use this to move back toward the home position, and when the cursor position is within 15% of the home-target distance (1.2 cm, Fig 2C), the cursor is shown again, to get back on to the home position.

In the rotated phase, these were used in a Process Dissociation Procedure (Jacoby, 1991; Werner et al., 2015) (”PDP”) in all three groups. In this Process Dissociation Procedure, people reach for a target without cursor, while either including or excluding their strategy (see below). The ’exclude’ strategy blocks are a measure of implicit adaptation, while the include strategy blocks should measure both implicit and explicit adaptation (Gastrock et al., 2020; Maresch et al., 2021b; Modchalingam et al., 2019; Werner et al., 2015). If implicit and explicit adaptation are additive, then subtracting reach deviations measured in include strategy blocks from reach deviations measured in exclude strategy blocks should be a measure of explicit adaptation. Whether or not this is true, if the responses are different in the two blocks, then participants can dissociate their strategic responses from unconscious adaptation, which shows that some degree of explicit adaptation is occurring. Note that if there is no difference in the responses, this is highly suggestive that no explicit adaptation is occurring, but not definitive proof.

### Aiming

Before training trials, participants in the aiming group would see the upcoming target and they could orient an arrow using the arrow keys on an extra key pad, to indicate the direction in which they would move their hand in order to get the cursor to the target. In the aiming group, the training blocks are called “Aim and reach” blocks, and the instructions they were given includes: “Point the arrow as accurately as you can to indicate the movement you want your hand to make, so that you can get the cursor on the target.”

### Instructions

#### Strategy

Participants in the instructed group received detailed instructions in between the aligned and rotated phase, while participants in the other groups had a break of about the same duration. The instruction included an animation of the perturbation, and explanation of a strategy to counter the perturbation (move in direction rotated 1 hour on a clock face). They were tested on understanding the strategy by drawing reaches on a clock face toward targets in all quadrants.

All groups were informed that the way the cursor moves changes in the next part of the experiment, and they were told that they have to figure out how to deal with this, and that they should remember what their strategy is, since they will be asked to reach while using this strategy and while not using this strategy, with this instruction:

”This is part 2 of the experiment. For these coming tasks, the filled cursor will move a bit differently, and you will need to compensate for this. However you compensate, keep that strategy in mind since you will be asked to use this strategy several times, when reaching without a cursor. Sometimes you will be asked to NOT use this strategy when reaching without a cursor.”

These instructions were read to participants before the rotated part of the task.

#### Process Dissociation Procedure

At the end of the rotated session, participants do no-cursor reaches in two kinds of blocks: either while including or excluding their strategy, with these instructions:

Include: “For THESE trials, make full use of any strategies you learned just now”.

Exclude: “For THESE trials, do not make use of any strategies you learned earlier and treat this as you did the original *’reach for target’* task in part 1”.

Importantly, the instructions about what to do in the PDP blocks were identical for all groups, such that they can not explain any differences in performance.

#### Procedure

All participants performed 264 reaching trials of various kinds (see Fig 2,3), organized into blocks of 8 trials. In each block, all 8 targets were used in a randomized order. The task started with an aligned phase with first 32 training trials, and 3 blocks of no-cursor trials (Fig 3 gray rectangles), with two blocks of training trials before the second and third no-cursor block. In the rotated phase, a rotation of 30° is applied to the cursors (Fig 3: black lines indicate perfect reaches). These included 96 training trials, followed by 3 sets of 2 no-cursor trial blocks. Adaptation is “topped up” before the second and third of these sets of no-cursor trials with 16 rotated training trials. The no-cursor trials in the rotated part are divided into 2 blocks: one where participants are instructed to include their strategy, and one where they do not include their strategy. In one group, the “aiming” group, every training trial is preceded by aiming: participants are shown the upcoming target and are asked to indicate where they will move their hand in order to get the cursor to the target. In the other groups, participants wait 1.5 s instead of aiming, to keep the total time roughly equal.

**Fig 3.**
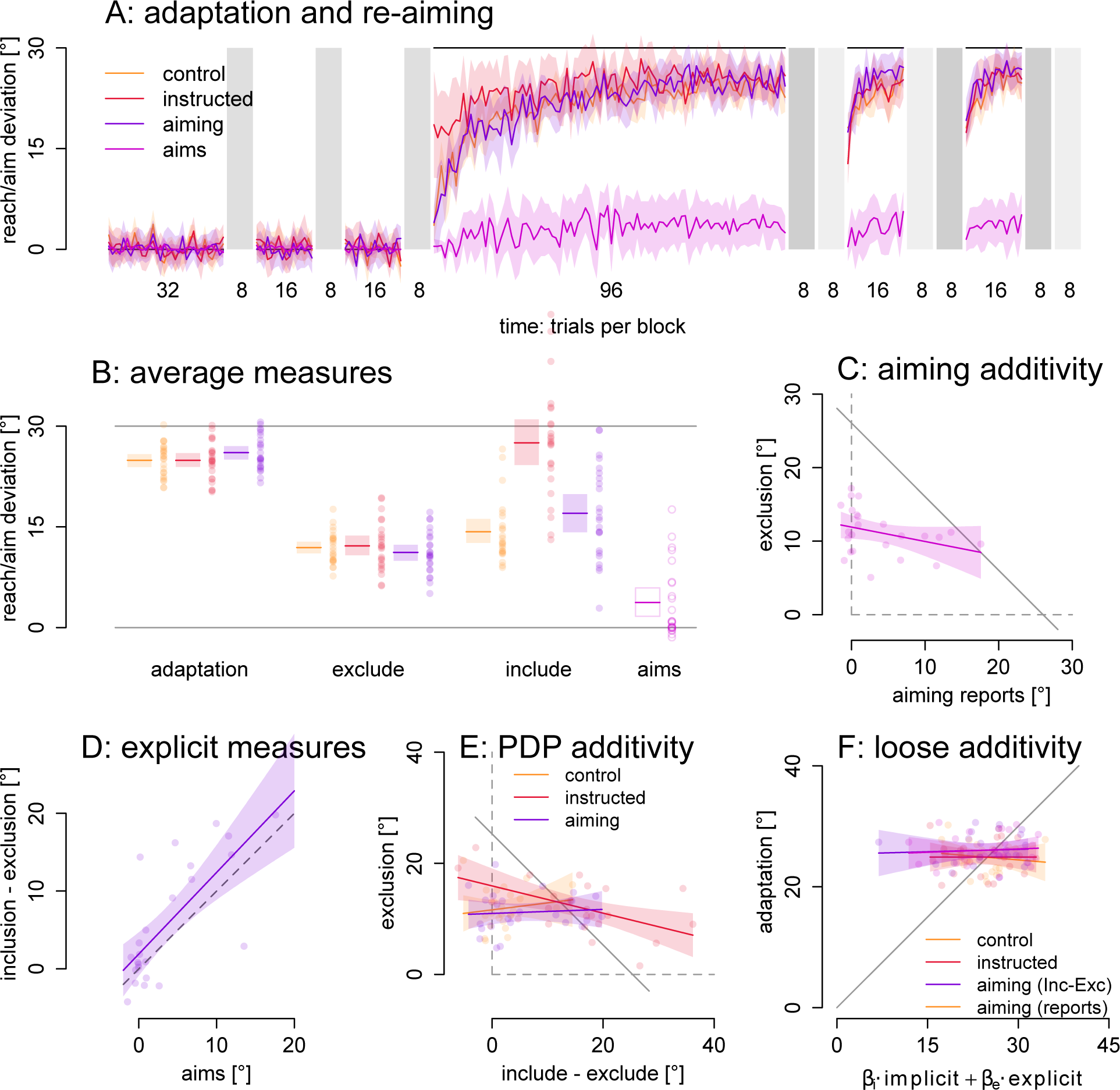
Adaptation measures and additivity. Areas in color denote 95% confidence intervals of the mean. Dots are individual participants. **A:** Adaptation and aiming responses throughout the perturbation schedule. No-cursor blocks in gray with inclusion and exclusion trials in the rotated phase. **B:** Averaged estimates of adaptation, exclusion and inclusion, and aiming reports. **C:** Strict additivity would predict that exclusion scores can be predicted from aiming scores and total adaptation and would lie close to the gray diagonal (slope -1), but there is close to no relationship between directions of aiming responses and exclude strategy reaches. **D:** Linear regression shows that aiming reports and include-exclude difference scores agree fairly well, such that we may be able to use include-exclude difference scores as a stand-in for aiming responses. **E:** Using indirect measures of explicit adaptation, we also can not confirm strict additivity, which would predict data lie close to the gray diagonal (with slope -1). **F:** Loose additivity would allow a weighted sum to predict total adaptation, with data over predictions on the gray unity line, but in this data set there is no relationship.

Participants were intermittently monitored during the experiment and encouraged to make straight, smooth reaches, to faithfully do no-cursor reaches according to instructions and to really consider where the aiming arrow should be pointed, if experimenters suspected the participant not to perform optimally.

#### Analyses

For all reaches (both training and no-cursor trials) we calculated the angular reach deviation at ⅓ the home-target distance (∼2.67 cm) and similarly used the aiming arrows angular deviation for comparison. We remove training reaches and re-aiming responses deviating more than 60° from the target (controls: 0, instructed: 1, aiming: 3, re-aiming responses: 0), and subsequently all angular deviations outside the average +/-3 standard deviation for each trial across participants within the group (controls: 15, instructed: 10, aiming: 12, re-aiming responses: 30). The maximum number of data points removed for any participant was 6 (control: 3, instructed: 4, aiming: 6, re-aiming responses 6), and many participants had no trials removed (control: 16, instructed: 19, aiming: 16, re-aiming responses: 12). No participants were removed, and no trials were removed from no-cursor data.

We use the average reach deviation in the last 8 trials before each of the no-cursor blocks, and subtracted the average in the aligned phase from the average in the rotated phase to assess final adaptation. In the aiming group we used the same blocks to assess final reaiming. We use all the include/exclude strategy no-cursor trials to assess implicit and explicit adaptation using the Process Dissociation Procedure. We subtract reach deviations from the aligned part of the task from the same measures obtained in the rotated part of the task to correct for baseline biases. This way we obtain adaptation in the training trials, with- and without strategy no-cursor reach deviations and re-aiming in the aiming trials. All of these are based on an equal number of trials, and taken from parts of the experiment that are as similar as possible.

We compare the exclusion reach deviations as a measure of implicit adaptation, and the final, total adaptation between the three groups. To assess explicit adaptation, we then compare the difference between inclusion and exclusion reach deviations. Within the aiming group we also compare the difference between inclusion and exclusion reach deviations, with the final aiming responses, as measures of explicit adaptation. All these allow testing of additivity on estimates of common measures of implicit and explicit adaptation processes.

### Strict Linear Additivity

We test additivity in 2 approaches. First, we investigate the notion total adaptation is simply the sum of explicit and implicit adaptation (Redding and Wallace, 1993) (cf. Fig 1):

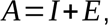

In particular, the way it is sometimes used to determine implicit adaptation:

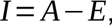

Which is equal to a linear model, expressing the relationship between implicit and explicit adaptation across participants (*p*):

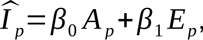

Where *β_0_ A* is the intercept, and *β_1_* is the slope (*β_0_*=1, *β_1_*=-1). It simply means that, with no explicit adaptation, implicit adaptation is equal to total adaptation (the intercept), and with every increase in explicit adaptation, there is an equally sized decrease in implicit adaptation (slope of -1; Fig 1D,E). This linear model directly follows from the additivity assumption, and would apply both within and across participants. A simulation to recover the slope of -1 from data generated according to this model plus noise, shows that the 95% confidence interval of the fitted slope includes -1 in ∼95% of simulations (Fig 1E). That is, if we decide that actual data is in accordance with additivity when the 95% confidence interval of the slope of a linear model predicting implicit from explicit adaptation includes -1, this is equivalent to a NHST statistic with ɑ=0.05. Hence, we will use this statistical approach.

### Loose Linear Additivity

To account for some systematic misestimates in either implicit or explicit adaptation, we use a second, less strict, model of additivity. Here, we predict total adaptation by summing *weighted* versions of implicit and explicit adaptation across all participants in each group:

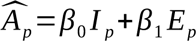

If this model of loose additivity works, then there should be a linear relation between predicted and actual adaptation with an intercept of 0 (no intercept) and a slope of 1. And here we test this by fitting a linear model on real adaptation over predicted adaptation, and we consider the model confirmed if the 95% confidence interval for the slope includes 1.

## Results

In our experiment, three groups of participants adapted to a 30° rotation: the main “aiming” group (N=24) that reported their aiming before every training trial by orienting an arrow in 1° steps, as well as an “instructed” group (N=24) that was made aware of the rotation and given a counter-strategy, and a “control” group (N=24). All groups repeated counterbalanced blocks of include- and exclude strategy no-cursor reaches (Werner et al., 2015) to assess implicit and explicit adaptation. Aiming reports are used as an independent measure of explicit adaptation. See methods for more details.

First, we find no group difference in adaptation (Fig 3A,B; F(2,69)=1.30, p=.28, η^2^=.036; BF10=0.316) nor in exclusion as measure of implicit adaptation (Fig 3B; F(2,69)=0.59, p=.555, η^2^=.017, BF10=0.185). Additivity applied to the Process Dissociation Procedure (PDP) measures would then predict equal inclusion deviations in the three groups (inclusion=implicit+explicit), but we find that the instructed group has higher inclusion deviations (F(2,69)=24.72, p<.001, η^2^=.417, BF10=1369327), with no difference between the other two groups. That is, the group averages of measures with built-in additivity do not seem to show additivity.

However, additivity should be tested by comparing the relationship between independent measures of explicit and implicit adaptation across various measures, which we do in the “aiming” group. We can see that the data does not appear to lie along the gray diagonal (Fig 3C) that depicts additivity based on the average total adaptation (cf. Fig 1D). Average adaptation in the aiming group is 26.0° (range: 21.5°-30.6°, sd: 2.6°). Aiming response range from -1.4° to 17.5° with an average of 3.7°, and exclude-strategy reach deviations range from 5.1° to 17.2° with an average of 11.2°. A linear regression of exclusion scores over aiming responses (Fig 3C; F(22,1)=2.8, p=0.11, R^2^ =0.073) has a slope of -0.196, and the 95% confidence interval of that slope (-0.437, 0.046) does not include -1, though it does include 0. That is, using independent measures of implicit and explicit adaptation, we can not find evidence for the additivity assumption.

Perhaps the difference between include and exclude strategy reach deviations are an acceptable stand-in for a measure of explicit aiming, such that we can test additivity in the other two groups. We test this in the aiming group, by predicting the inclusion-exclusion difference scores from the reported aims, with a linear regression. The two measures largely agree (Fig 3D; F(1,22)=25.84, p<.001, R^2^=0.5192, intercept=1.9, slope=1.05, slope 95% CI: 0.62 - 1.47) — in this data set, but not everywhere (Heirani Moghaddam et al., 2021; Maresch et al., 2021b). Given this, we also test strict additivity in the three groups using the inclusion-exclusion differences as a measure of explicit adaptation (Fig 3E), but find that in none of the three groups the 95% confidence interval of the slope includes -1 (aiming: -0.17 - 0.24, control: -0.26 - 0.51, instructed: -0.41 - -0.08). This result does rely on the additivity assumption, and yet it shows no relationship between implicit and explicit adaptation. However, the finding based on aiming reports uses independent measures and should be considered more reliable.

### Do the fast and slow processes map onto measures of explicit and implicit adaptation?

Next, we test a particular application of the additivity assumption, in a common state-space model with two adaptive processes. In this model, the so-called “slow” and “fast” processes are strictly additive (Smith et al., 2006), and are sometimes assumed to map onto implicit and explicit adaptation respectively (McDougle et al., 2015). In previous work (McDougle et al., 2015), the similarity between averaged aiming reports and the fast process fitted to group data is striking. However, given that the model also assumes additivity and that the measure of explicit adaptation matches the model’s fast process fairly well, it has to be the case that the subtractive estimate of implicit adaptation matches the slow process equally well. That is, while the literature supports a link between the model’s fast process and an independent measure of explicit adaptation, the link between the slow process and implicit adaptation is fully dependent on the largely untested assumption of strict linear additivity, and should not be relied upon, indiscriminately. We test this here in the data from the aiming group in the experiment described above.

## Methods

### Two-Rate Model

Using data from the experiment described above, we test a state-space model with a strong additivity assumption (Smith et al., 2006), and two learning processes that are sometimes assumed to map onto implicit and explicit adaptation (McDougle et al., 2015). The model posits that motor output on trial t (Xt) is the straightforward, unweighted sum of the states of a slow and fast learning process:

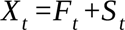

Where the states of both the fast and slow process are each determined by a learning rate (L) applied to the error on the previous trial, and a retention rate (R) applied to the state of each process on the previous trial:

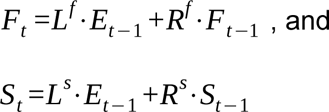

This is further constrained by *_Ls <Lf_*, and *_Rf <Rs_*.

### Parameter Recovery Simulation

The model typically is fit on data with a different perturbation schedule, but the 2 blocks of no-cursor reaches causing some decay in learning, followed by relearning, may suffice for a reasonable group-level fit. To test this, we ran a parameter recovery simulation for the model using the training reach deviations in the control group (fit model 1000 times to data simulated as original model + random noise drawn from a normal distribution with mean=0 and a standard deviation equal to the square root of the MSE of the original model fit). This shows that the parameters can be recovered quite well (not shown, bootstrapped data on OSF: https://osf.io/9gpfk). For each parameter the originally fitted value falls in the 95% interval of the recovered values (L^s^: 0.028 < 0.035 < 0.043; L^f^: 0.087 < 0.105 < 0.129; R^s^: 0.991 < 0.994 < 0.996; R^f^: 0.832 < 0.885 < 0.912). This means that fitting the model to reach deviations in this perturbation schedule is robust, at the group level (Albert and Shadmehr, 2018), and we can use the model to test if its processes map onto the measures of explicit and implicit adaptation throughout learning.

### Aware and Unaware Learners

On average, we find only a small re-aiming strategy in the aiming group (Fig 3B: 3.74°; t(23)=3.38, p=.001, η^2^=0.497). However, we observed that the distribution of the differences between inclusion and exclusion reach deviations for participants in the aiming group is bimodal (relative log-likelihood based on AICs of unimodal distribution compared to bimodal: p>0.001). The two peaks are close to the average inclusion-exclusion differences in the control and in the instruction group (Fig 4A-C). This likely indicates that for 9/24 participants, the addition of aiming had the same effect as instructions, but no effect for the remaining 15 participants.

**Fig 4.**
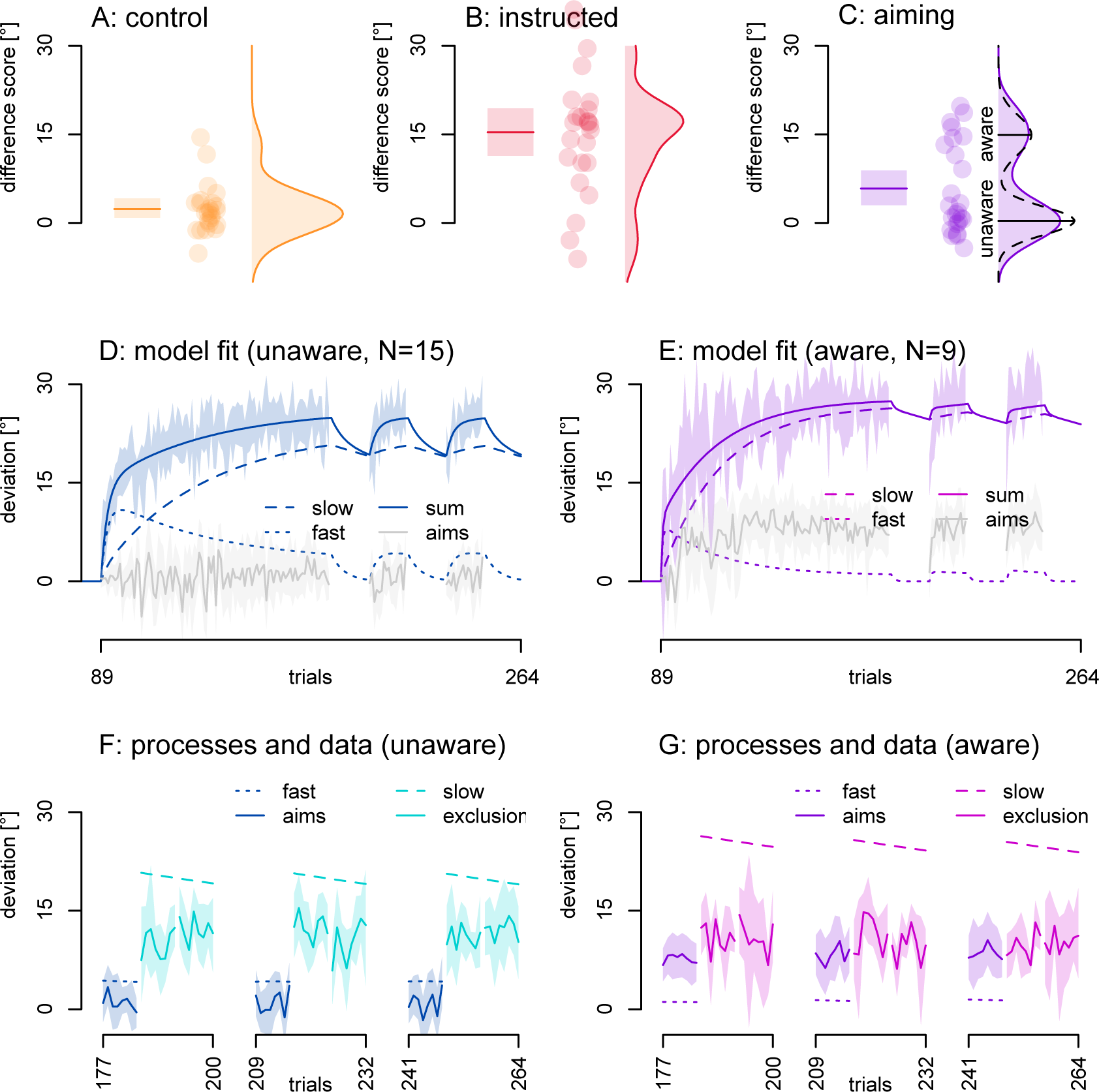
Additive state-space model. **A-C:** Distributions of differences between include- and exclude strategy reach deviations, depicted as density plots. Dots represent individual participants, solid lines and shaded regions in bar plots represent the mean and 95% confidence interval. **C:** In the aiming condition, the distribution is bi-modal (p<.001) suggesting aware and unaware subgroups. The underlying processes in the two-rate model fits should differ between these two subgroups if the fast process maps onto explicit adaptation. Thus, we fit the model separately for each subgroup (D-E). **D & E:** Shaded area represents the 95% confidence interval of reach deviations and the solid line the two-rate model fitted to these reach deviations, for the unaware aimers (D) and the aware aimers (E). **F:** Two-rate model fits. In the aware subgroup, the fast process (smooth continuous purple line) for the most part is outside the 95% confidence interval of the aiming responses (jagged continuous purple line with 95% confidence interval). In the unaware subgroup, the confidence interval of the aiming responses (jagged continuous dark blue line with 95% confidence interval) is near zero and excludes the fast process on most trials. Similarly, the slow process of each subgroup’s model fit (the dashed lines) are above the 95% confidence interval for the exclude strategy reach deviations (jagged continuous lines in the no-cursor blocks: light blue for the unaware aiming group, light pink for the aware aiming group). Thus, in both subgroups, the fast process does not predict aiming responses, and the slow process does not predict strategy exclusion reach deviations.

We split the aiming group into aware aimers and unaware aimers, as these subgroups should have different levels of explicit adaptation. We fit a separate two-rate model to the averaged training reach deviations in the aware aimers as well as those in the unaware aimers. We then test whether or not the fast process falls in the 95% confidence interval of each subgroups’ aiming responses, and whether or not the slow process falls in the 95% confidence interval of exclusion reach deviations in each subgroup. The comparison between the slow process and exclude-strategy reach deviations (implicit) can only be done in the no-cursor blocks, when the processes have presumably saturated. The comparison between aiming responses and the fast process is then also done when the processes have saturated: during the 3 blocks of 8 trials just before the no-cursor blocks. If the model processes fall within the 95% confidence interval of the data, this would support additivity as implemented in the two-rate model.

## Results

We test if the state-space model’s slow process maps onto exclusion trials (implicit) and if the fast process maps onto aiming responses (explicit). We separately fit the state-space model to the averaged reach deviations in aware and unaware aimers (Fig 4C). In the aware aimers (Fig4D, purple), the fast process (purple smooth continuous line) is lower than aiming reports (purple jagged line with 95% confidence interval), and vice-versa in the unaware aimers (dark blue). In both subgroups the 95% confidence interval of the aiming data excludes the fitted fast process (except for a few trials at the start). In both subgroups the slow process (dashed lines) is higher than exclusion trials during all no-cursor blocks (95% CIs of data exclude model processes; pink=aware aimers, light blue=unaware aimers). confirming that the slow process does not capture implicit adaptation (Bansal et al., 2022; Ruttle et al., 2021). That is: the strictly additive processes of the state-space model do not align with direct measures of implicit and explicit adaptation in this data set.

### Testing additivity across the field

While it stands to reason that adaptation is larger when both implicit and explicit adaptation processes contribute (Benson et al., 2011; Mazzoni and Krakauer, 2006; Neville and Cressman, 2018), few studies indicate some form of additivity (Bond and Taylor, 2015; Sülzenbrück and Heuer, 2009). Other studies don’t show additivity (Gastrock et al., 2020; Modchalingam et al., 2019; Schween et al., 2018; Werner et al., 2015) or they show partial additivity, or additivity only in some participants (Bromberg et al., 2019; Neville and Cressman, 2018) or it is unclear to us whether or not the data support additivity (Heirani Moghaddam et al., 2021; Maresch et al., 2021b). So far, we rejected additivity based on one dataset only. It would be easy to dismiss these findings as a one-time occurrence. However, if additivity does not hold, this has implications for the field. Hence, we test strict and loose linear additivity in a number of other data sets, with independent if not simultaneous measures of explicit, implicit and total adaptation. We collected data sets from previous work, both from other labs and our own, to test strict and loose additivity beyond the single experiment described above with data representing various approaches in the field.

## Methods

We either asked the original authors (Bond and Taylor, 2015; Brudner et al., 2016; Neville and Cressman, 2018; Taylor et al., 2014) or downloaded data from public repositories (Decarie and Cressman, 2022; Maresch et al., 2021b; Modchalingam et al., 2022, 2019), and used one unpublished data set sent to us by Dr. Jordan Taylor as well as one incomplete data set from a study conducted in our lab (D’Amario et al., 2024). This is not an exhaustive literature search, but should represent several types of equipment, approaches and research groups, and it should increase power. See Table 1 for more information.

**Table 1:** Sources of external data. Number of trials are indicated, measures are from trials when adaptation has saturated and measures should be stable, so they can be averaged for more precision. The exceptions are the data sets from the papers in the first row, where measures of implicit adaptation are taken during its washout. Here we fit an exponential function to the first 16 trials, and use the starting value as an estimate of implicit adaptation.

| Study | Data source | Adaptation | Implicit | Explicit |
| --- | --- | --- | --- | --- |
| Sources with independent aiming and exclusion |  |  |  |  |
| Bond & Taylor (2015); Taylor et al. (2014); Brudner et al. (2016); unpublished data set | private exchange | last 16 training trials | starting value of an exponential function fitted to the first 16 washout trials | last 16 aiming reports of training |
| Schween et al. (2018 exp. 1 & 2 ; 2019 exp. 1, 2 & 3) | private exchange | average across 9 targets | average across targets straddling the averaged generalization curve's peak | average across 9 targets |
| D'Amario et al. (2024) | In-lab exchange | 16 training trials | 16 exclusion trials | 8 aiming trials |
| Sources with PDP difference scores |  |  |  |  |
| Neville & Cressman (2018) | private exchange | last 9 training trials | 9 exclusion trials in in last pre-pause PDP block | 9 inclusion - 9 exclusion trials in last pre-pause PDP block |
| Modchalingam et al. (2019) | OSF repository linked in paper: https://osf.io/mx5u2/ | last 18 trials of the first rotated training | 4 x 9 exclusion trials | 4 x 9 inclusion - 4 x 9 exclusion trials |
| Maresch et al. (2020) | OSF repository linked in paper: https://osf.io/6yj3u/ | Last 8 trials in "top-up" block | Last 16 exclusion trials | Last 16 inclusion - last 16 exclusion trials |
| Decarie & Cressman (2022) | Supplementary file with paper: https://link.springer.com/article/10.1007/s00221-022-06352-4 | Last 15 trials of rotated reach training | last 6 exclusion trials | last 6 inclusion - last 6 exclusion trials |
| Modchalingam et al. (2023) | https://osf.io/a7k6s/ | 2 x 9 training trials before PDP blocks | 2 x 9 exclusion trials | 2 x 9 inclusion - 2 x 9 exclusion trials |

### Normalization

As before, we test if the confidence intervals for the slopes of the strict and loose models of linear additivity include -1 and 1 respectively, both on all individual data sets, and all the data combined. The rotations varied between 15° and 90°, so we need to normalize the measures to be able to compare data across groups. Since the additivity assumption predicts adaptation (not rotation), we normalize relative to each participants’ estimate of total adaptation. If additivity holds, the normalized measures of implicit and explicit adaptation should then sum to 1. We do test normalization by rotation on all data combined as well, and this yields highly similar results (Fig 5, last two rows). However, to stay more in line with the tested additivity assumption, we use only the adaptation-normalized scores for the further exploration of aggregate data.

**Fig 5.**
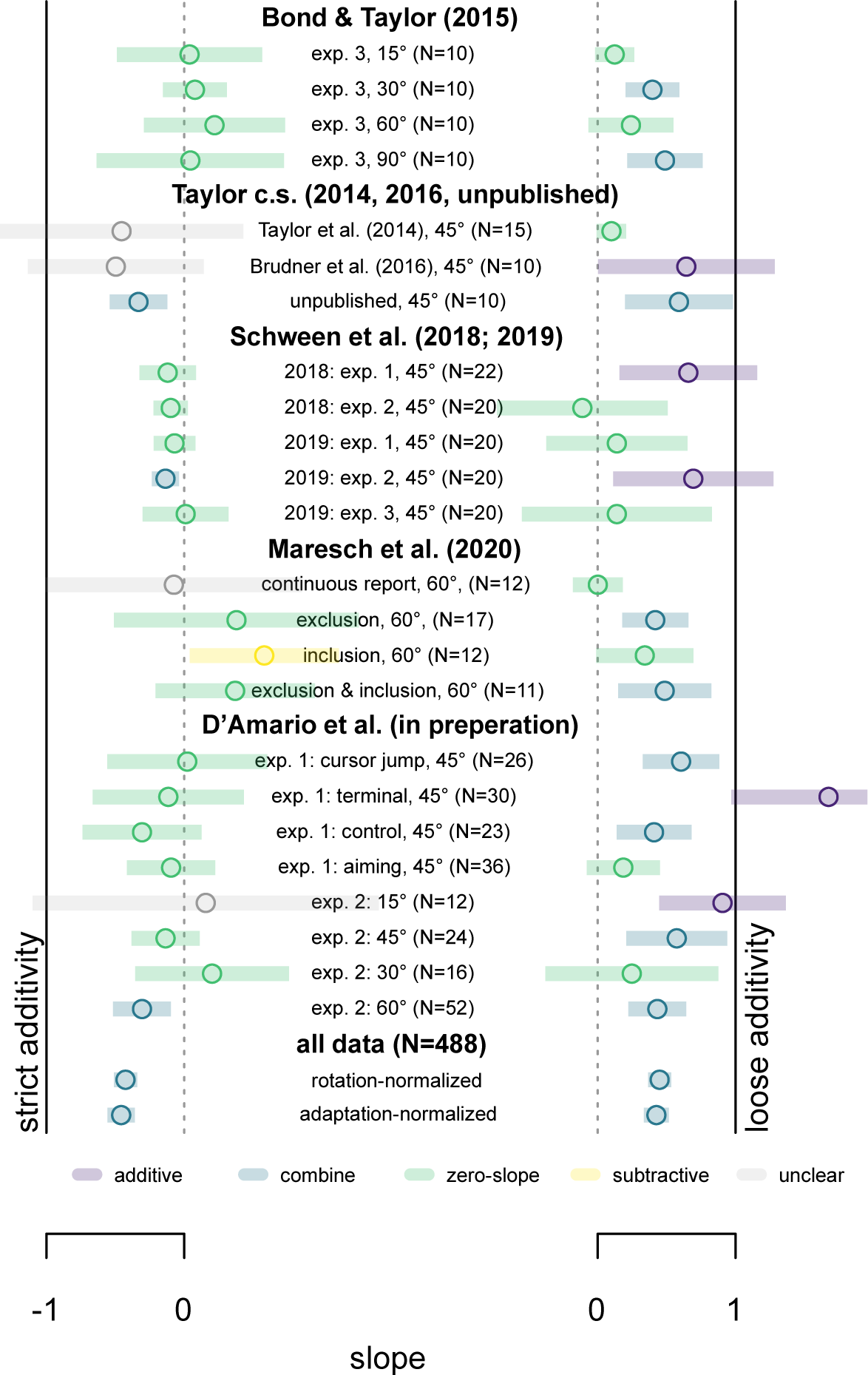
Additivity in a cross-section of studies with independent measures of implicit and explicit adaptation. In each group (rows) we test strict additivity (left) by testing if -1 (continuous black line) is included in the 95% confidence interval (colored bars) of the slope of a linear model fitted to independent measures of both implicit and explicit adaptation. No groups demonstrate strict additivity. We also test loose additivity (right), where in each data set a free weight is set to both implicit and explicit contributions before summing them to predict total adaptation. Here the 95% confidence interval (colored bars) of the slope of predicted over actual adaptation should include 1 (continuous black line). Only 5 of 24 subsets show loose additivity. In the combined data (bottom two rows), there does seem to be support for some form of combining explicit and implicit adaptation.

## Results

The literature is divided on the additivity assumption of implicit and explicit adaptation (Maresch et al., 2021a). That’s why we re-analyzed data from 11 studies (Bond and Taylor, 2015; Brudner et al., 2016; D’Amario et al., 2024; Decarie and Cressman, 2022; Modchalingam et al., 2022, 2019; Neville and Cressman, 2018; Schween et al., 2019, 2018; Taylor et al., 2014). There are 24 groups (N=488) with independent measures of implicit and explicit adaptation, and 16 groups (N=325) using the Process Dissociation Procedure, that has an independent measure of implicit adaptation, but not of explicit adaptation. For every participant we took an average across multiple trials at the end of training to estimate explicit, implicit and total adaptation when each of these measures should have saturated (the “stepwise” participants (Modchalingam et al., 2022) are assessed 4 times, at 4 different rotation sizes, for 43 ’groups’ total).

First, we look at data with truly independent measures of explicit and implicit adaptation (top rows of table 1; Fig 5). We divide each participants’ estimates by their total adaptation, and for each subgroup accept strict additivity (Fig 5: left) or loose additivity (Fig 5: right) if the 95% CI for the slope of the model includes -1 or 1 respectively (as above; Fig 1D&E, Fig 3D&E).

None of the groups support strict additivity (see Fig 5), and only 5 of 24 subgroups support loose additivity. Consequently, in all participants combined (Fig 5, bottom two rows) there is no support for additivity either.

We also test the relationship between estimates of implicit and explicit adaptation, when these are not strictly independent (bottom rows of table 1; Fig 6). Here, this coincides with studies using the Process Dissociation Procedure only (or some variant), where explicit adaptation is taken to be the difference between inclusion and exclusion trials, while implicit adaptation is independently estimated by exclusion trials. That is, the additivity assumption is built into the estimate of explicit adaptation. Despite this, however, none of the 19 groups support strict additivity, and only one supports loose additivity.

**Fig 6.**
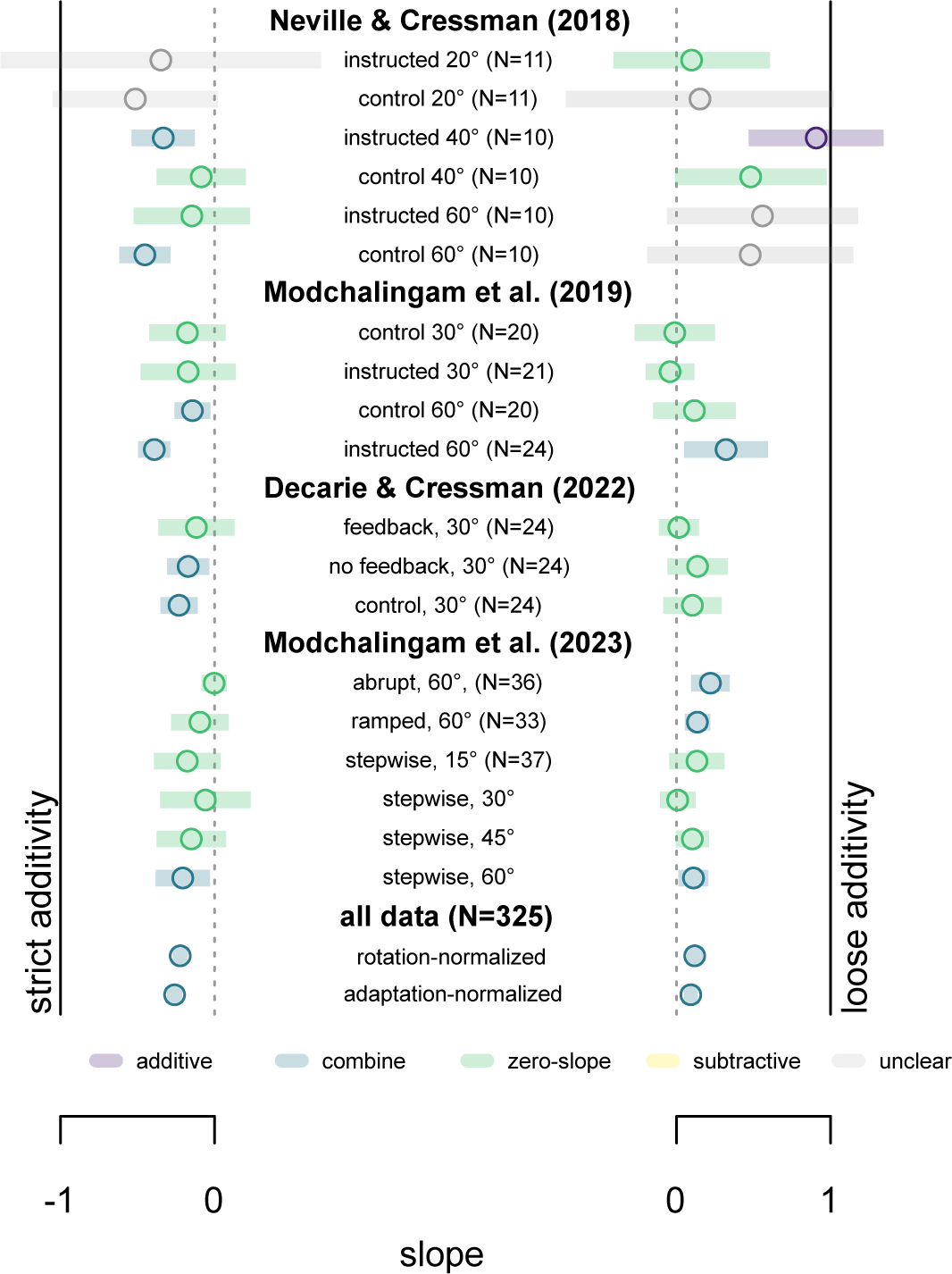
Additivity in a cross-section of studies where implicit and explicit adaptation are not independent. All data comes from studies using Process Dissociation Procedure. Here total adaptation is not relied on, as in most studies using measures of aiming, but the measures of explicit and implicit adaptation are related (see methods). Strict additivity (left) is supported if the 95% confidence interval (colored bars) of the slopes fitted to implicit over explicit adaptation include -1 (continuous black line). Loose additivity (right) is supported if the 95% confidence interval (colored bars) of a line fitted to the predicted over the measured adaptation includes 1 (continuous black line). No groups support strict additivity, and only 1 of 19 support loose additivity. The overall relationship between implicit and explicit adaptation (bottom rows) is weaker than in studies with independent measures (Fig 5), despite the measures used here relying on a form of additivity.

### Maximum Likelihood Estimation of adaptation

The aggregate data does show a combined effect: slopes *are* different from 0 (Fig 5, 6 and 7A). If contributions from implicit and explicit adaptation are not added, the question is: how are they combined? We test one possible answer, inspired by maximum likelihood estimation, using the collection of data sets from above. In this approach adaptation is estimated by weighting explicit and implicit adaptation by the inverse of their variance. That is: the more reliable adaptation process contributes most to total adaptation. Similar approaches provide powerful explanations for combined sensory estimates, so this may be a more biologically plausible mechanism.

**Fig 7.**
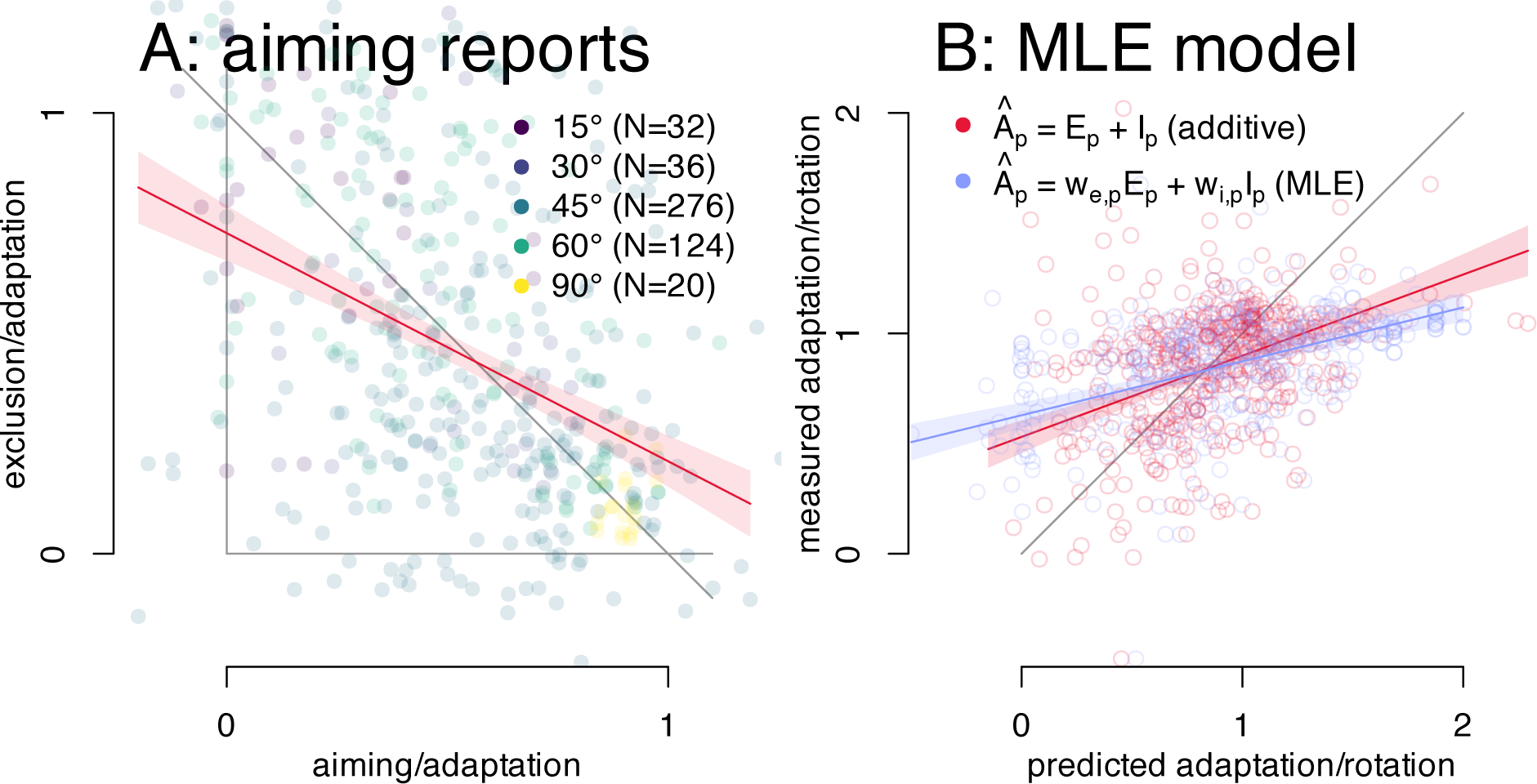
Maximum Likelihood Estimate combining implicit and explicit adaptation. **A:** Adaptation-normalized data and a linear regression of implicit over explicit adaptation (N=488). The gray diagonal indicates perfect additivity. **B:** Predicted adaptation based on strict additivity and on Maximum Likelihood Estimates for 387 participants. The gray diagonal indicates perfect prediction.

## Methods

### Maximum Likelihood Estimates

To test if implicit and explicit adaptation are combined based on their relative reliability, with a greater contribution for the more reliable process, we applied a maximum likelihood estimate. Here each of the processes is weighted by the relative inverse of their variance within each participant:

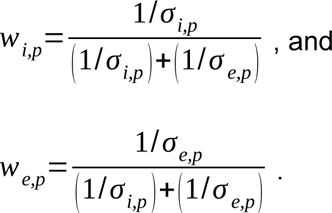

Where *w_i,p_* and *w_e,p_* are the weights for implicit and explicit adaptation respectively for participant *p*, and *σ_i,p_* and *σ _e,p_* are the variance of implicit and explicit measures for participant *p*. These are based on the data, but require a direct measure of each. This is only possible for independent measures of implicit adaptation (such as exclusion trials) as well as for explicit adaptation (such as re-aiming) with a fair number of individual trials available. This restricts us to data from 4 papers (Bond and Taylor, 2015; Brudner et al., 2016; Maresch et al., 2021a; Taylor et al., 2014) plus one unpublished group, and the incomplete data set from our lab, for 488 participants adapting to rotations of 15°, 30°, 45°, 60° and 90° (see Fig 7A). To account for the different rotation sizes, we use rotation-normalized estimates of implicit, explicit and total adaptation. The estimates of total adaptation for each participant *p* are then:

*A_p_ =w_i,p_ E_p_ +w_e,p_ I_p_*, in the maximum likelihood estimate, and simply

*A_p_ =E _p_ +I_p_*, for the comparable additive model.

To be clear, in each case *A_p_* is the predicted adaptation for participant *p*, based on: *E_p_* and

*I _p_*, the average measure of explicit and implicit adaptation for the same participant.

The weights in maximum likelihood estimates sum to 1, but in the strictly additive approach they would sum to 2. We also tested a sum of weights of 2, but this results in a slightly worse fit. We tested some other variants as well: a freely fitted sum of weights, adding a constant, or scaling the contributions from implicit and explicit adaptation independently. These all produce comparable results, so we opted for presenting the simplest model.

## Results

Some models of adaptation use Bayesian mechanisms to weigh different contributions (Berniker and Kording, 2011; Tsay et al., 2022), and these mechanisms are not linearly additive, so here we test one as an alternative way explicit and implicit adaptation are combined. In particular we test a Maximum Likelihood Estimate (Ernst and Banks, 2002) (MLE). The weight applied to the average of each contribution is the relative inverse of its’ respective variance for every participant which means that a separate estimate of variance of both implicit and explicit adaptation is required (details in methods). This is only possible in 488 participants, and we test how well a strictly additive and an MLE mechanism predict the measured adaptation. As can be seen (Fig 7B), neither predicts adaptation very well, and the maximum likelihood approach does slightly worse than the additive approach. I.e., maximum likelihood estimation is an unlikely alternative to strict additivity. The way implicit and explicit contributions are combined in motor adaptation remains unclear.

## Discussion

In this study we tested whether measures of implicit and explicit adaptation linearly add in total adaptation. In particular, we tested the validity of deriving an indirect, “quasi” measurement of implicit adaptation by subtracting a measure of explicit adaptation from a measure of total adaptation. It seems clear that explicit and implicit processes interact to determine the amount of total adaptation, but we find no evidence supporting linear addition. Below, we discuss what this means for our understanding of explicit and implicit adaptation, implications for future practice, as well as the interpretation of some previous findings.

While both implicit and explicit processes contribute to adaptation, and a plethora of work has been done testing what either process is sensitive to, there are two main sources of problems with how we understand the contribution of each. First, the methods we use to measure both implicit and explicit adaptation all have their issues (Maresch et al., 2021b). The second source is the focus here; we do not know the neural mechanism by which contributions from different learning processes are combined in behavior.

In our experiment, as well as in a cross-section of other data, there was unexpectedly little evidence of an additive relationship between independent measures of explicit and implicit adaptation. This means that implicit adaptation can not be estimated by subtracting explicit adaptation from total adaptation, or vice versa. Previous work on visuomotor adaptation might need to be re-evaluated if their findings on implicit adaptation rely on a strict additivity assumption, e.g. when subtracting a measure of explicit adaptation from total adaptation to get a measure of implicit adaptation (usually in studies relying on measures of re-aiming). Conversely, explicit adaptation is sometimes gauged by subtracting reach deviations in exclusion trials from those in inclusion trials (as we did here). This is done in studies relying on the Process Dissociation Procedure or PDP, although this appears less common. Nevertheless, it should be clear that this approach suffers from the same lack of support as more popular subtractive measures. In the absence of a known mechanism for combining implicit and explicit adaptation either approach may be justified in some cases. Perhaps one of these subtractive measures is slightly better than the other depending on whether the study is primarily focussed on explicit or implicit adaptation. However, neither should not be relied on indiscriminately and a better alternative is needed.

Most current models of adaptation combine separate (neural) processes by adding their outputs. Here we chose the two-rate model (Smith et al., 2006) as an example since it is well-known and yet still relatively straightforward. While the original study publishing the model convincingly shows there are multiple processes at play in adaptation, the mechanism used to combine the output of separate processes in this model is not necessarily neurally plausible. Crucially, this is not limited to this model, but extends to any model using linear addition of neural processes. Neural net-based models would be an exception. Regardless, not only are the terms in our equations important, so are the operators. That is; while using addition is a good simplification while modeling, we should keep in mind that it is a simplification that does not necessarily reflect actual neural processes.

Some models explore new directions however, suggesting that separate adaptation processes can be sub-additive (Albert et al., 2022), which could correspond to “loose” additivity here, depending on the fitted weights. Others use Bayesian mechanisms (Berniker and Kording, 2011; Tsay et al., 2022), such as Maximum Likelihood Estimates, that are well-known in the literature on multi-modal perception (Ernst and Banks, 2002). However, neither of these can explain the data we considered here.

In some experiments or measures this study relies on, the magnitude of either process could have been systematically misestimated. For example, some decay might occur between regular reach trials and whenever a measure of implicit adaptation is taken, resulting in under-estimation. Or perhaps, between reaching with and without a strategy, the strategy is not fully disengaged resulting in over-estimation of implicit adaptation. Alternatively, if participants only train with a single target, the largest reach aftereffects (or strategies) don’t necessarily occur at the trained target, so that tests at the trained target would underestimate implicit (or explicit) adaptation. There might also be other reasons for systematic under- or overestimating of either implicit or explicit adaptation. And at first glance, the aggregate data (Fig 7A) indicates a linear relationship with systematically underestimated implicit adaptation. We have tried to overcome this with our “loose” additivity approach. Here a weight is applied to both estimates of implicit and explicit adaptation before they are added as a prediction of total adaptation. While “loose” additivity fared somewhat better in some cases (Fig 5, 6), it does not seem that systematic under-estimation (or over estimation) for any reason can explain why strict additivity does not hold up in the current study.

The re-analysis of aggregate data from across the field shows a weak relationship between measures of implicit and explicit adaptation. And while we didn’t see any relationship in our own data set, it seems more plausible that the two interact in overall behavior. What then, do we know about how implicit and explicit processes combine? First, given the lack of clear patterns, either the measures (Maresch et al., 2021b) or the underlying implicit and explicit adaptation processes could be highly variable and may saturate (Kim et al., 2018) - or both. Second, the mechanism by which the two processes combine is unknown, but not strictly additive. Third, we should consider that this relationship is likely influenced by other factors such as interactions between processes (Albert et al., 2022; Tsay et al., 2022), task contexts (Bond and Taylor, 2015; Decarie and Cressman, 2022; Gastrock et al., 2020; Modchalingam et al., 2019) or goals or motivations of participants. Beyond motor adaptation, these notions may apply to other types of learning as well.

What we learn from this study can be summed up in two recommendations. First, in order to investigate implicit and explicit processes, we should use independent measures of both. Currently, it seems that the best candidates are re-aiming responses for explicit, and strategy exclusion trials for implicit adaptation, although it is also unknown how neural adaptation processes map onto these behavioral measures. Second, the field should try to understand the (neural) mechanism by which various adaptation processes are combined to shape behavior.

## Acknowledgements

We are grateful to Dr. Jana Maresch and MSc. Amelia Decarie for making their data publicly available, and to Dr. Erin Cressman, Dr. Jordan Taylor, and Dr. Raphael Schween for sharing some of their data with us. We are also thankful for feedback from Dr. Opher Donchin, Dr. Gunnar Blohm and Dr. Scott Albert, an informal review by Dr. Nina van Mastrigt and Dr. Jeroen Smeets as well as anonymous reviews from eNeuro that all shaped the content of this paper.

